# Feature-specific patterns of attention and functional connectivity in human visual cortex

**DOI:** 10.1101/869552

**Authors:** Kirstie Wailes-Newson, Antony B Morland, Richard J. W. Vernon, Alex R. Wade

## Abstract

Attending to different features of a scene can alter the responses of neurons in early- and mid- level visual areas but the nature of this change depends on both the (top down) attentional task and the (bottom up) visual stimulus. One outstanding question is the spatial scale at which cortex is modulated by attention to low-level stimulus features such as shape, contrast and orientation. It is unclear whether the recruitment of neurons to particular tasks occurs at an area level or at the level of intra-areal sub-populations, or whether the critical factor is a change in the way that areas communicate with each other. Here we use functional magnetic resonance imaging (fMRI) and psychophysics, to ask how areas known to be involved in processing different visual features (orientation, contrast and shape) are modulated as participants switch between tasks based on those features while the visual stimulus itself is effectively constant. At a univariate level, we find almost no feature-specific bottom-up or top-down responses in the areas we examine. However, multivariate analyses reveal a complex pattern of voxel-level modulation driven by attentional task. Connectivity analyses also demonstrate flexible and selective patterns of connectivity between early visual areas as a function of attentional focus. Overall, we find that attention alters the sensitivity and connectivity of neuronal subpopulations within individual early visual areas but, surprisingly, not the univariate response amplitudes of the areas themselves.

## 1.1 Introduction

Attention directed toward spatial locations or visual features can influence both behaviour and physiology. In general, attention tends to increase the relative sensitivity of neurons representing the attended location or feature (e.g. Duncan and Humphreys, 1989; Corbetta et al., 1990; Luck et al., 1997; Reynolds et al., 2000; Martínez-Trujillo and Treue, 2002; Serences and Boynton, 2007), rendering participants faster and more sensitive when detecting changes at attended locations, or in pre-cued features (Posner et al., 1980; Posner and Cohen, 1984; Corbetta et al., 1990).

Given many early visual areas have been identified as having intrinsic preferences for particular features (for a review, see Kanwisher, 2010), it might seem obvious that attention towards those features would modulate activity within entire areas. For areas with high specificity for a single feature, this does seem to be the case (e.g. (Corbetta et al., 1991). fMRI BOLD responses in hMT+ increase during attention toward a visual motion stimulus (O’Craven et al., 1997; Huk et al., 2001). Increased hV4 activation has been identified in response to attention to chromatic stimuli (Chawla et al., 1999; Liu et al., 2003; Schoenfeld et al., 2007) and attention to faces and places can drive robust responses in the fusiform face area (FFA) and parahippocampal place areas (PPA) respectively (O’Craven et al., 1999).

However, recent work suggests that attention acts primarily through a gain control mechanism, whose primary effect is to alter inter-neuronal noise correlation (Cohen and Maunsell, 2009). If this is correct, we might expect to see relatively little attentionally-driven change in overall activity in earlier visual areas. While individual neuronal sub-populations might become more or less correlated in their trial-to-trial responses, normalization would serve to reduce any long-term response differences (Verhoef and Maunsell, 2017).

This hypothesis is consistent with two findings from the literature: first that early visual areas exhibit relatively little overall change in activity when subjects switch low-level visual tasks (Kamitani and Tong, 2006; Sumner et al., 2008; Brouwer and Heeger, 2009; Seymour et al., 2009, 2010; Song et al., 2011). Secondly, that attentional modulation is detected readily by EEG, which is sensitive to the level of correlated noise in large-scale neuronal populations (Martinez et al., 2001; Di Russo et al., 2003; Wang and Wade, 2011; Verghese et al., 2012).

If attention drives changes in the relative sensitivity and noise correlation of neuronal sub-populations, it should alter activity at the level of feature maps within each visual area. These changes might be detected using multivariate fMRI methods, which in the past, have enabled researchers to decode participants’ featural attentional focus from fMRI BOLD activity (Kamitani and Tong, 2005, 2006; Sumner et al., 2008; Brouwer and Heeger, 2009; Freeman et al., 2011; Song et al., 2011).

Finally, attention can change patterns of activity *across* areas as well as within a single region. Such changes could act at the level of input or output layers, altering not just the sensitivity of neurons performing within-area computations, but also the type of information passed between areas. In support of this hypothesis, it is clear that brain-wide changes in functional connectivity are associated with different mental states (Greicius et al., 2003; Fox et al., 2005) and, more specifically, previous literature has identified attentionally-driven changes in patterns of connectivity amongst regions involved in visuospatial tasks (Fox et al., 2005; Gao et al., 2013; Spadone et al., 2015).

Here, we looked for evidence of differential *univariate* or *multivariate* responses in regions-of-interest (ROIs) within the visual cortex, as well as differences in inter-areal *connectivity*. In the multivariate case, we also asked if successful classification was driven by coarse-scale topographical maps or fine-scale voxel-level sensitivities.

Surprisingly, we found very few univariate differences in the regions we examined. However, voxel-level analyses revealed above-chance decoding of attentional focus in *all* visual ROIs, with successful decoding driven by fine-grain participant-specific differences in voxel activity. Connectivity analyses revealed greater connectivity between visual regions during passive viewing than during feature-specific directed attentional focus, and these feature-specific connectivity patterns changed as a function of the ROI subset examined.

## 1.2 Materials and methods

### 1.2.1 Participants

12 participants were recruited (8 female, mean age 25 years). All participants had normal or corrected-to-normal vision and provided informed consent. Ethical approval for the study was granted by the University of York Department of Psychology and York Neuroimaging Centre ethics boards. Participants completed two 1.5-hour scanning sessions as part of this experiment, during which we collected high resolution anatomical scans, population receptive field mapping (pRF) and attentional modulation data.

### 1.2.2 Experimental Design

#### 1.2.2.1 Behavioural Psychophysics

To control for difficulty and levels of attention, prior to fMRI scanning, each participant completed 1 hour of psychophysical testing, to identify their 75% correct detection thresholds for each visual feature (orientation, contrast and shape). Prior testing revealed no significant differences in difficulty (indexed by loglinear d’) or reaction time across the attentional focus conditions F(2,22) = 0.61, *p* = 0.553, F(2,22) = 1.36, *p* = 0.275 (no significant Bonferroni-corrected post-hoc tests) respectively. This ensured task difficulty, and associated attentional effort was constant across attention conditions within our fMRI experiment.

Stimuli were presented on a ViewPixx monitor (120Hz, 1920×1220 pixels resolution) at 57cm viewing distance. Stimulus presentation was performed on a Shuttle XPC SZ87RG high-end graphics system with an Intel Core i7-4790K processor at 4GHz and a NVIDIA GeForce GTX970 graphics card with 4GB DDR5 memory. All stimuli and experimental procedures were controlled by Psychtoolbox 3.0.12 routines (Brainard, 1997; Pelli, 1997).

To estimate detection thresholds, we used a Bayesian adaptive staircase design (QUEST) (Watson and Pelli, 1983). Feature-specific initial estimates of threshold (0.3 radians orientation, 30% contrast, 0.08 radial amplitude modulation/shape) were provided with 0.5-unit standard deviation. Our stimuli were variants of a radial frequency pattern (RFPs): radially modulated circular contours (Wilson and Wilkinson, 1997; Wilkinson et al., 1998; Ivanov and Mullen, 2012). This stimulus allowed for an investigation of shape processing and reflected naturalistic stimuli to a greater extent than traditional Gabors (Lawrence et al., 2016; Vernon et al., 2016). Reference RF pattern stimuli had a 2.0° average radius, 0° orientation and 50% contrast (see Figure.1A).

**Figure.1:**
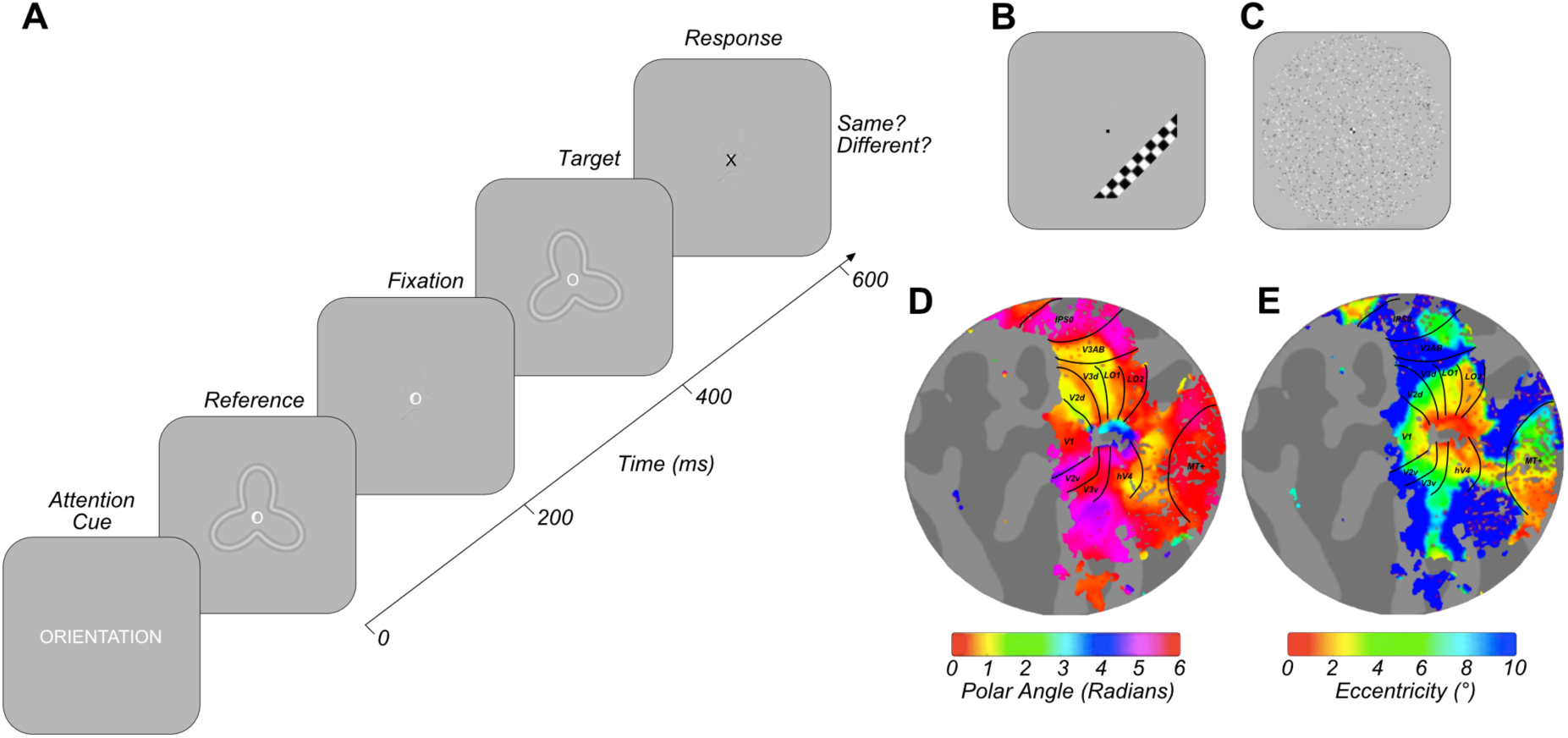
Exemplar stimuli and retinotopic maps. A) Example psychophysics/fMRI trial structure B) Example high-contrast phase-reversing checkerboard drifting bar stimulus used to gain estimates of population receptive field size, as in Dumoulin & Wandell (2008). C) Full-field motion stimulus used to help delineate MT+ ROIs, (adapted from Huk et al., 2002; Fischer et al., 2012; Maloney et al., 2013). D & E) Exemplar left hemisphere retinotopic maps with ROI border overlays presented on flattened cortical representations for one participant. Visual ROIs were defined on the basis of eccentricity (A) and polar angle (B) in accordance with the processes described in Dumoulin, Wandell & Brewer (2007).

Attended features were cued by a central white fixation letter present across trials (‘O - orientation’, ‘C - contrast’ or ‘S - shape’). Trials began with a 200ms presentation of a grayscale reference stimulus against a mean luminance background, followed by a 200ms inter-stimulus interval with presentation of the letter-fixation cue only. A target stimulus was then presented for 200ms, with a change in the attended visual feature derived from the initial estimate and individual participant’s previous trial performance. For each feature, a stimulus change could be in one of two directions (with equal probability) - clockwise versus anticlockwise orientation, high contrast versus low contrast, ‘spiky’ versus ‘smooth’ shape. This was followed by a central black fixation cross for 800ms, during which participants made indicated the perceived direction of change between the reference and target stimulus (‘A’ or ‘L’ keyboard press). Participants were informed via a toned ‘beep’ if their response was correct or incorrect. Each participant completed 50 trials (alongside 10 discarded practice trials at the start of each run) to provide an estimated 75% correct detection threshold for each feature. Each feature-specific staircase was repeated three times with a participant’s final detection thresholds reflecting the average of these three repetitions. Each run lasted approximately two minutes.

#### 1.2.2.2 fMRI Attentional Modulation

The fMRI experimental design followed a similar procedure to the psychophysics paradigm. However, here we wished to examine the effects of participants directing attention toward changes in individual visual features. To ensure participants maintained a constant attentional load, they constantly monitored the stimulus for the a near-threshold-level change in an attended feature (ignoring changes in uncued features). In this respect, our fMRI experiment differed from the psychophysics-each visual feature was not modulated on every trial, and changes in multiple visual features could co-occur. The fMRI experiment was also designed such that we presented the same average stimulus (i.e. the probability of a change in each feature was constant across blocks), so that any effects we identify should be driven by differing attentional focus as oppose to stimulus-driven effects. The timing of each trial within our fMRI experiment, however, was identical to the psychophysical testing paradigm.

Prior to each attentional focus block a 1.5s visual cue directing participants attentional focus was presented (white text; ‘ORIENTATION’, ‘CONTRAST’, ‘SHAPE’ or ‘PASSIVE’) against a mean luminance background. During passive blocks, participants were instructed to view the stimulus without directed attention and were not required to respond. As before, each trial began with a 200ms presentation of the reference RFP with a white central fixation letter matching the cued feature of attention (‘O’, ‘C’, ‘S’ or ‘P’), followed by a 200ms presentation of the fixation cue. This was followed by a 200ms presentation of the target RFP stimulus, which could vary in no, singular or multiple visual features with respect to the reference stimulus. The degree of change between the reference and target RFP was double the participant’s previously-recorded feature-specific 75% averaged correct detection threshold. Each feature varied in only one direction (anticlockwise orientation, high contrast and spikier shape). For each feature, the target stimulus differed from the reference on approximately 20% of trials, hence a constant level of visual modulation was present across blocks on average. A central fixation letter was then presented for 800ms, during which participants made a 2AFC (same/different) response. Each trial was 1.5s duration, with a block consisting of 10 trials (15s). Each block was followed by a black central fixation cross (7.5s) allowing BOLD signal to return to baseline. Blocks were presented in a pseudo-randomised order, with the randomised four-block cycle presented four times within each run.

### 1.2.3 Functional Neuroimaging

#### 1.2.3.1 fMRI data acquisition

Visual stimuli were presented using a PROpixx DLP LED projector (Vpixx Technologies Inc., Saint-Bruno-de-Montarvile, QC, Canada) with a long throw lens which projected the image through a waveguide behind the scanner bore and onto an acrylic screen. Presented images had 1920×1080 pixels resolution and 120Hz refresh rate. Participants viewed the stimulus at 57cm viewing distance within the scanner.

A Shuttle XPC SZ87RG high-end graphics system with Intel Core i7-4790K processor at 4GHz and a NVIDIA GeForce GTX970 graphics card with 4GB DDR5 memory matched to the system used in the Psychophysical testing were used to control the fMRI experiment. All stimuli and experimental procedures were controlled by MATLAB 8.5.0 (2016a) in conjunction within Psychtoolbox 3.0.12 routines (Brainard, 1997; Pelli, 1997). During scanning behavioural responses and scanner trigger pulses were acquired using a fibre-optic response pad Forp-932 (Current Designs, Philadelphia, PA).

fMRI data were collected at the York Neuroimaging Centre using a GE 3T Excite MRI scanner (GE Healthcare, Milwaukee, WI). Structural scans were obtained using an 8-channel head coil (MRI Devices Corporation, Waukesha, WI) to minimise magnetic field inhomogeneity. Population receptive field and attentional modulation data were obtained with a 16-channel posterior head coil (Nova Medical, Wilmington, WI) to improve signal-to-noise in the occipital lobe. Two high-resolution, T1-weighed full-brain anatomical structural scans were acquired for each participant (TR 7.8ms, TE 3.0ms, T1 450ms, voxel size 1 × 1 × 1.3mm^3^).

pRF scan sessions consisted of 6.5-minute stimulus presentation runs collected using a standard EPI sequence (TR 3000ms, TE 30ms, voxel size 2.0 × 2.0 × 2.5mm^3^, flip angle 20°, matrix size 96 × 96 × 39mm^3^). pRF parameters were obtained using procedures very similar to those described by Dumoulin and Wandell (Dumoulin and Wandell, 2008) (see Figure.1B).

Attentional modulation scan sessions consisted of an average of 6, 6:42-minute runs, containing 134 volumes of data, including 3 dummy TRs later removed to allow for the scanner magnetisation to reach a steady-state. 39 slices were acquired in a bottom-up interleaved acquisition order (TR 3000ms, TE 30ms, field-of-view 19.2cm^3^, matrix size = 96 × 96 mm, voxel size 2.0 × 2.0 × 2.5mm^3^, flip angle 90°).

During the pRF and attentional modulation scanning sessions, one 16 channel coil T1-weighted structural scan with the same spatial prescriptions as the functional scans was acquired to aid in the alignment of the functional data to the T1-weighted anatomical structural scan (TR 2100ms, TE 8.6ms, field-of-view 19.2cm^3^, matrix size 512 × 512mm, voxel size 0.38 × 0.38 × 2.5mm^3^, flip angle 90°, 39 slices).

#### 1.2.3.2 fMRI Pre-processing

To improve grey-white matter contrast, the two T1 high-resolution anatomical scans were aligned and averaged together using the FLIRT FSL tool (Jenkinson et al., 2012). This averaged T1 was automatically segmented using a combination of FreeSurfer (Fischl, 2012) and FSL, and manual corrections were made to the segmentation using ITK-SNAP (Teo et al., 1997).

Functional data were analysed using MATLAB 2016a (Mathworks, MA) and VISTA software (Vista Lab, Stanford University). Between- and within-scan motion correction was performed to compensate for motion artefacts occurring during the scan session. Any scans with > 3mm movement were removed from further analysis (no attentional modulation runs were removed on the basis of excessive movement). The Vista rxAlign tool was then used to co-register the 16-channel coil T1-weighted structural scan to the 8-channel coil T1-weighted full-brain anatomical scan. We applied a manual alignment by using landmark points to bring the two volumes into approximate register, followed by a robust EM-based registration algorithm to fine-tune the alignment (Nestares and Heeger, 2000). The final alignment was visually inspected to ensure the automatic registration procedure optimised the fit. This alignment was then used as a reference to align the functional data to the full-brain anatomical scan. These functional data were then interpolated to the anatomical segmentation.

#### 1.2.3.3 Population receptive field mapping

To probe attentional modulation across the visual system, we focused our analysis on early visual regions with clear feature-specific preferences or organisation related to our stimulus modulations. The discrete cytoarchitecture of V1, consisting of regular blobs and interblobs with differential spatial and contrast tuning might result distinct activation patterns associated with orientation and contrast. (Horton and Hubel, 1981; Livingstone and Hubel, 1988; Song et al., 2011). Regions LO-1 and LO-2 are clearly-defined, retinotopic areas on the lateral surface of visual cortex (Larsson and Heeger, 2006) that have been identified as having a particular significance in both shape and orientation processing (Larsson and Heeger, 2006; Silson et al., 2013). We also selected two additional visual ROIs, a ‘ventral’ area, hV4 and a ‘dorsal’ area V3A/B; regions with no clear expected patterns of attentional modulation to summarise patterns of attentional modulation across the visual cortex. For example, hV4 has been implicated in contrast, orientation and shape processing (Ghose and Ts’ O, 1997; Dumoulin and Hess, 2007; Sani et al., 2013).

We additionally noted attentional modulation driven by changes in cued task demands must involve feedback signals from higher cortical areas such as the IPS to lower-level visual regions (e.g. Di Russo et al., 2003; Bressler et al., 2008; Lauritzen et al., 2009). We therefore included the IPS as a separate ROI of particular interest in our analysis of connectivity.

pRF parameters (eccentricity, polar angle and size) were estimated for each voxel using the standard pRF model within mrVista (Dumoulin and Wandell, 2008). Following Wandell et al., (2007) we manually delineated 9 bilateral ROIs: V1, V2, V3, V3A/B, hV4, LO-1, LO-2, MT+ and IPS0 on cortical flat maps on the basis of polar angle reversals and eccentricity for each participant (see Figure.1D and Figure.1E).

Seven participants possessed previously-collected MT+ localiser data (the design adapted from (Huk et al., 2002; Fischer et al., 2011; Maloney et al., 2013), defining MT+ on the basis of responses to motion versus static stimuli (see Figure.1C). For these participants, we used these motion-defined MT+ ROIs. To ensure consistency and examine reliability across these different MT+ identification techniques, each participants’ structural space was transformed to Talairach coordinates using the seven landmarks outlined in (Ryu et al., 2010). A reference spherical 8mm MT+ ROI was created from standardised Talairach coordinates centred on [LH: -47 -76 2, RH: 44 -67 0] (Dumoulin et al., 2000). All defined-MT+ ROIs overlapped with the standardised control MT+ spherical ROIs. Using the same process, a 5mm spherical control ROI of the primary auditory cortex (A1) was created for each participant with standardised coordinates centred on [LH: -49 -20 9, RH: 48 -21 10] (Lacadie et al., 2008).

#### 1.2.3.4 Attentional Modulation

General linear models (GLM) were implemented to test the contribution of stimulus condition to the BOLD time course (Friston et al., 1998). The default two-gamma Boynton HRF from the SPM8 toolbox was used (Penny et al., 2006) and we fit the model to an averaged time course of BOLD signal changed for each stimulus condition by minimising the sum of squared errors between the predicted timeseries and the measured BOLD response.

The first GLM analysed bottom-up stimulus feature change events. Events were classified according to the nature of the stimulus change occurring within a 1 TR (3s) period, regardless of attentional focus (orientation, contrast, shape, no change and multiple change events). Multiple change events reflected two different feature stimulus changes occurring within a single TR. Feature change events (orientation, contrast and shape) reflected when both trials within a TR contained a change in the same feature (e.g. orientation change, orientation change), or a feature change and no feature change (e.g. orientation change, no change). This resulted in 52 to 141 events per stimulus change condition. The second GLM analysed the contribution of attentional focus (15s blocks of directed attention to orientation, contrast, shape or passive viewing). This resulted in a vector of 24 average beta weight estimates for each voxel at the multivariate level.

### 1.2.4 Statistical analysis

#### 1.2.4.1 Univariate Analyses

Feature-specific attentional modulation univariate beta weights (orientation, contrast and shape) were averaged (per participant) and compared with the passive beta-weight through Wilcoxon signed-rank tests for each ROI. Independently for event and attentional modulation analyses, feature-specific univariate betas were also analysed through one-way repeated measures ANOVAs to assess stimulus-driven and attentional modulation differences in BOLD signal modulation across orientation, contrast and shape conditions.

#### 1.2.4.2 Multivariate Analyses

In each ROI we selected the 100 voxels that explained the largest amount of variance across conditions. These multivariate beta weights were z-scored across voxels and used as input to a support vector machine (SVM) (‘LIBSVM’ toolbox optimised for Matlab with a radial basis function) (Chang and Lin, 2011) to decode either bottom-up stimulus change or featural attentional focus in two separate analyses using leave-one-out cross validation for each participant. We first assessed multi-class decoding accuracy, supplying orientation, contrast and shape data simultaneously using the ‘one-against-one’ approach to produce a single classification accuracy score for each participant (Knerr et al., 1990). For the attentional modulation data, pairwise classification was also performed assessing classification accuracy for each combination of attentional conditions (orientation versus contrast, orientation versus shape, contrast versus shape) to identify the driving forces behind successful multi-class decoding. Classification accuracies were then assessed against chance performance through one-sample Wilcoxon signed-rank tests for each ROI.

To investigate the spatial localisation of feature-specific attentional modulation across voxels, additional pairwise SVM classification was performed between each attentional focus condition and the passive viewing data. The weighted mean of support vectors from these classifications was calculated to provide a feature-specific attention ‘preference’ for each voxel within an ROI. These voxel ‘preference’ weights were back-projected onto an interpolated polar grid (6° eccentricity, 360° polar angle, across 500 samples) to reflect voxel activation as a function of each voxels preferred visual angle and eccentricity, extracted from pRF analysis. Voxel preference for each visual feature was thresholded at +/−1.7 z-score (*p* <.05) and displayed to indicate the spatial distribution of attentional modulation. We averaged data over eccentricities between 1.5° and 3.5° of visual angle and plot the resulting average activation as a function of polar angle to provide an intuitive summary of spatial attentional focus. For reference, a standard RF pattern (0° orientation, 0.2 radial amplitude modulation) was overlaid. This back-projection analysis was performed for V1, V2, V3 and hV4 ROIs, as other ROIs lack a high-resolution representation of the full visual field.

#### 1.2.4.3 Timeseries analyses

To quantify functional connectivity between ROIs, participant-specific multivariate timeseries data were extracted (grouping TRs by attentional focus condition) and underwent noise removal (fit and removal of a grand mean) to eliminate scan-to-scan differences in raw amplitude intensity. These multivariate data were then averaged across all voxels within an ROI to provide a single univariate timeseries for each attention condition. Non-parametric Kendall’s tau correlations were performed for all pairwise combinations of ROI (V1, V3A/B, hV4, LO-1, LO-2 and IPS0) – generating a correlation matrix for each condition. To assess the overall similarity of connectivity patterns, the correlation matrix for each attentional focus condition was itself converted to a vector (removing self-to-self correlations) and normalised via participant-specific global mean extraction and Fisher-z transformation. The average (Fisher-z transformed) correlation coefficient was then computed to provide a singular number summarising group connectivity within each attentional task. The difference in correlations between conditions was analysed via a one-way repeated measures ANOVA with Bonferroni-corrected post-hoc tests.

Additionally, we investigated ROI-specific *connectivity patterns* as a function of attentional condition. For each of the ROIs; V1, V3A/B, hV4, LO-1, LO-2 and IPS0, we extracted the data reflecting the correlation of this ROI with all others (e.g. V1-V3A/B, V1-hV4, V1-LO-1, V1-LO-2, V1-IPS0) for every participant. We then sampled data from all participants (12 samples with replacement) with a sample reflecting a full complement of ROI-specific correlation data for each attentional focus condition and calculated the mean across these samples. To simulate noise/chance data in this analysis, we took the same 12 samples selected with replacement for each condition, and for each pairwise comparison of conditions, we switched the condition labels approximately 50% of the time, keeping ROI-ROI relationships constant and calculated the average across these scrambled condition-specific datasets. For both the observed and noise data, we calculated the root mean squared error (RMSE) distance between each pairwise combination of condition vectors as a measure of difference in patterns of connectivity across the ROI profile of interest between different attentional modulation conditions. This process was repeated across 10,000 iterations for each ROI comparison (5 comparisons). Across all iterations, we then calculated the percentage of observed RMSEs for a pairwise comparison falling below the RMSE of the comparable simulated noise distribution. Any percentile below 5% indicates a difference in ROI-specific patterns of connectivity between two attentional modulation conditions significantly larger than predicted by chance (*p* <0.05).

## 1.3 Results

### 1.3.1 Univariate analysis: stimulus-driven events

We first asked whether our near-threshold stimulus modulations altered the BOLD signal at a univariate level. The one-sample Wilcoxon-signed rank test of average BOLD signal across orientation, contrast and shape conditions was significantly different from zero in almost all ROIs examined (Benjamini-Hochberg adjusted p-values reported) (V1: Z(11) = 2.27, *p* = 0.034, V3A/B: Z(11) = 2.35, *p* = 0.034, LO-1: Z(11) = 3.06, *p* = 0.009, LO-2: Z(11) = 2.98, *p* = 0.009, A1: Z(11) = -2.12, *p* = 0.041). The only exception was hV4 where averaged bottom-up stimulus-driven activity was not significantly different from zero (Z(11) = 1.88, *p* = 0.060). Overall, we find that our subtle stimulus modulations do generate a small but significant change in BOLD contrast in most visual areas while auditory cortex ROI (A1) demonstrated small but significant *negative* BOLD responses on average.

We then asked whether this BOLD modulation changed depending on the type of stimulus change. For example, BOLD signal changes in LO-1 and LO-2 have been associated with bottom-up changes in orientation and shape respectively (Larsson and Heeger, 2006; Silson et al., 2013) although the stimulus manipulations in these reports were far larger than those used in this study.

One-way repeated measures ANOVAs Benjamini-Hochberg corrected for the number of ROIs revealed no significant differences in univariate BOLD modulation between stimulus-driven changes in orientation, contrast and shape in any visual ROI (*p* > 0.05) (see Figure.2B). Negative BOLD responses were found across all three stimulus-driven change conditions in A1, again with no significant differences related to particular stimulus manipulations.

**Figure.2:**
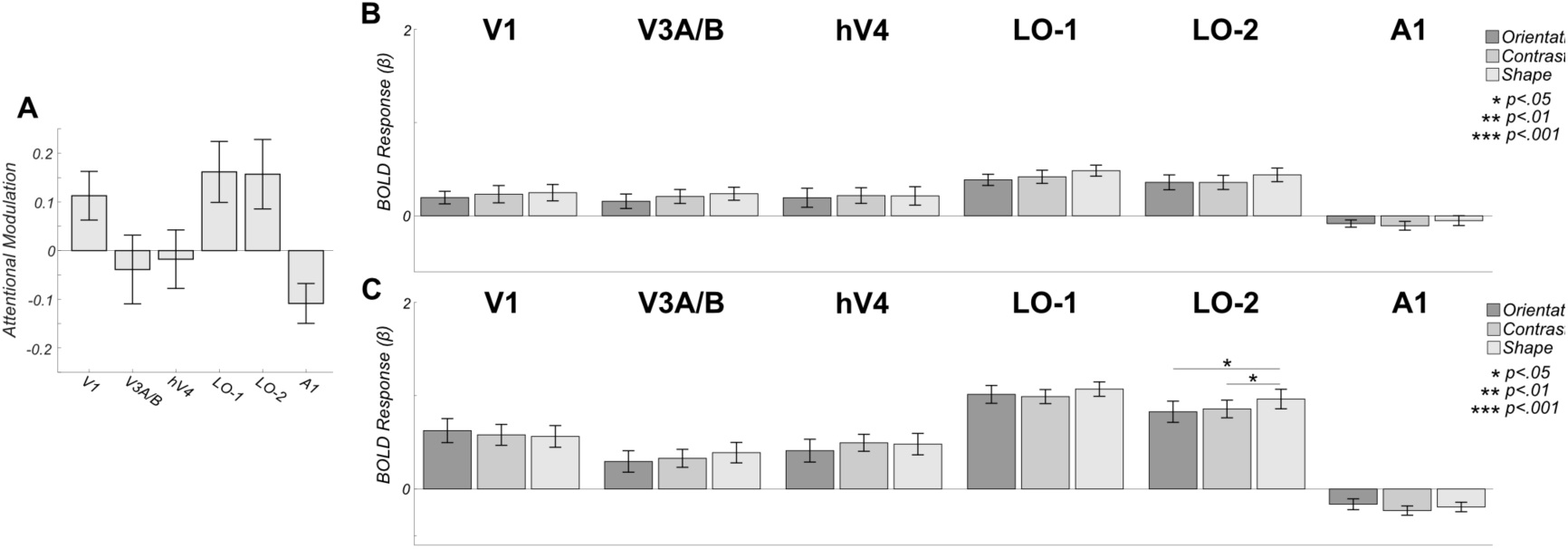
Univariate responses to both attention and stimulus modulation are weak. A) Mean BOLD modulation due to attention: positive values reflect greater BOLD amplitude during directed attention (averaged across attentional conditions). Data are shown for 5 visual ROIs: V1, V3A/B, hV4, LO-1 and LO-2 as well as a control area (auditory cortex-A1). No areas exhibit significantly larger BOLD modulation the attend vs passive comparison. B) Mean bottom-up stimulus responses to individual modulation events averaged over all attentional conditions. No areas exhibit differential responses to bottom-up stimulus modulations. C) Feature-specific attentional modulations averaged over all modulation types. Area LO-2 shows a slight increase in BOLD amplitude when subjects attend to shape. All error bars are +/− 1 SEM and significance asterisks indicate Benjamini-Hochberg corrected values.

### 1.3.2 Univariate analysis: attentional modulation

In the absence of differential BOLD responses relating to stimulus change, we looked for evidence of BOLD modulation as a function of attentional task. Although the majority of visual ROIs examined (excluding V3A/B and hV4) exhibited, on average, greater positive BOLD modulation during directed attention versus passive viewing, Wilcoxon-signed rank tests revealed these trends were not significant (adjusted *p* > 0.05) (see Figure.2A). A1 demonstrated a trend towards greater positive modulation during passive viewing versus averaged directed attention, but again this was not significant (adjusted *p* > 0.05).

Although on average we found no evidence of attentionally-driven BOLD modulation, it is possible that such changes were present for individual attentional condition types. We therefore conducted one-way repeated measures ANOVAs, Benjamini-Hochberg corrected for the number of ROIs, which revealed no significant differences in attentional modulation across the three featural attentional focus conditions (orientation, contrast and shape) within across almost all visual ROIs examined (*p* > 0.05) (see Figure.2C). There was a single exception to this null finding: Within LO-2, Bonferroni-corrected post-hoc tests revealed significantly greater attentional modulation for attention to orientation compared to either shape (F(2,22) = 6.61, *p* = 0.034, post-hoc *p* = 0.011) or contrast (*p* = 0.026). Again, no significant differences in attentional modulation were evident within A1 (F(2,22) = 3.29, *p* = 0.112).

Broadly, our univariate analysis showed that neither the bottom-up stimulus manipulations nor the top-down attentional demands had strong *differential* effects on the BOLD signal in early visual cortex. Area LO-2 was an exception, demonstrating a weak but significant differential response for attention to orientation compared to the other task conditions.

### 1.3.3 Multivariate Analysis: pattern classification

The stimulus modulations that subjects detected were extremely subtle. Nevertheless, as shown in Figure.2B, we detected BOLD activity with an amplitude significantly greater than zero time-locked to these modulations. We therefore asked if we could decode the identity of these bottom-up responses based on the pattern of responses they elicited in each ROI. To do this, we performed a three-way multivariate pattern classification analysis on the event-related responses, simultaneously classifying orientation, contrast and shape modulations. One-sample Wilcoxon signed-rank tests versus chance (33.33%) Benjamini-Hochberg corrected for the number of ROIs (6), revealed that the type of *bottom-up* stimulus driven modulations could not be decoded at rates significantly above chance in any ROI examined (*p* >0.05). Mean decoding accuracy ranged from 31.81 to 34.18% (see Figure.3A).

**Figure.3:**
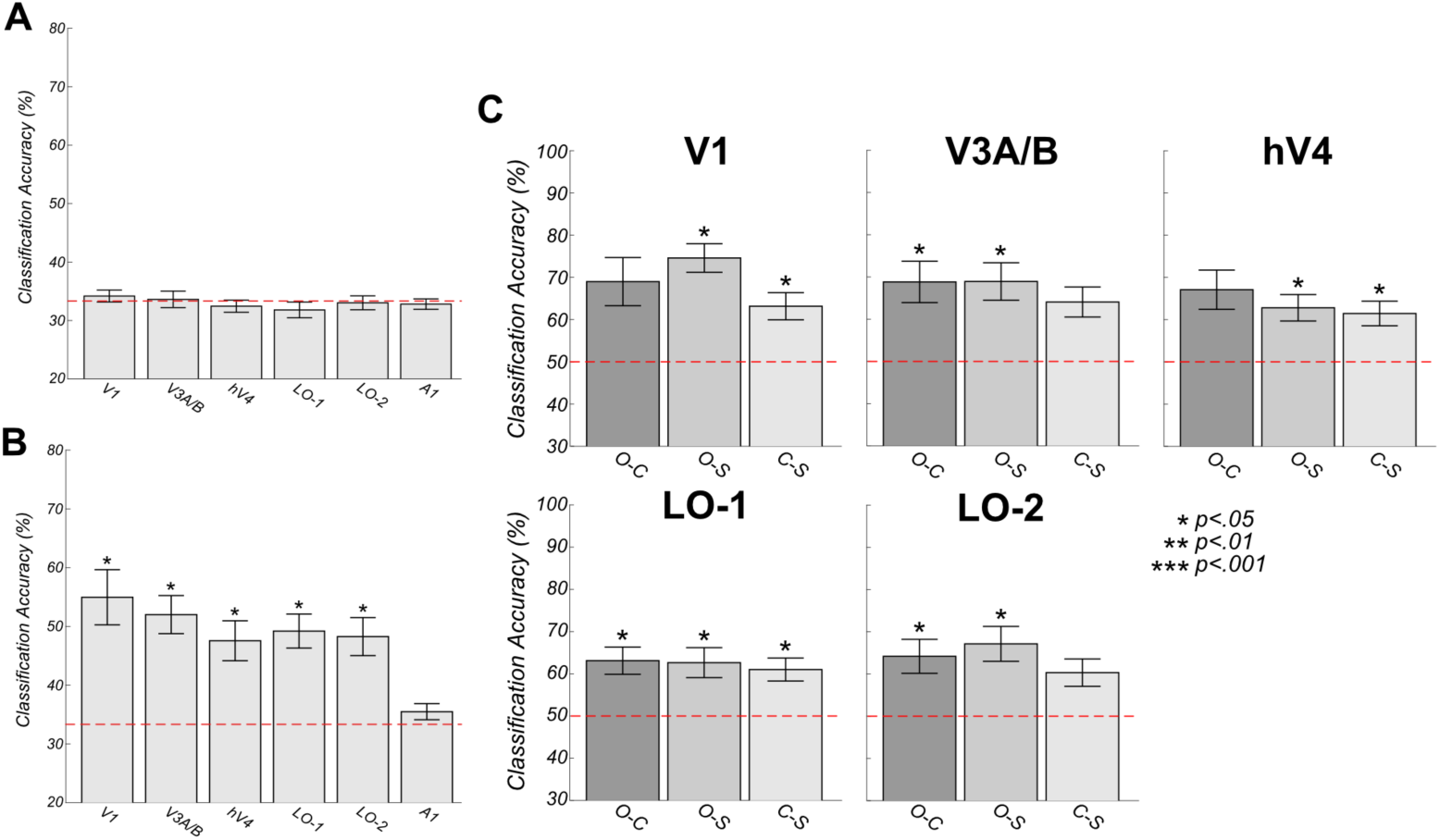
Multivariate Support Vector Machine Decoding. Voxel-level responses with individual ROIs are modulated by attentional state, but not by bottom-up changes in stimulus features. A) Overall three-way decoding accuracies. Stimulus change cannot be accurately decoded in an ROI examined. B) Attentional state can be decoded above chance in all ROIs except A1. C) Two-way classification accuracies across pairwise combinations of attentional focus (orientation versus shape, orientation versus contrast and contrast versus shape). Voxel patterns in all areas differ significantly between attention to orientation and shape. Error bars reflect +/− 1 SEM and significance asterisks indicate Benjamini-Hochberg corrected values.

The overall BOLD activity linked to attentional modulation was also significantly greater than zero (Figure.2C). However, in comparison to the bottom-up results, multivariate pattern classification of attentional state achieved accuracies significantly greater than chance in all visual ROIs examined (V1: Z(11) = 2.90, *p* = 0.014, V3A/B: Z(11) = 3.06, *p* = 0.014, hV4: Z(11) = 2.75, *p* = 0.018, LO-1: Z(11) = 3.06, *p* = 0.014, LO-2: Z(11) = 2.94, *p* = 0.014) (see Figure.3B). Mean classification accuracy ranged from 35.49 to 54.96% across ROIs. As expected, classification was not significantly different from chance within the auditory control ROI A1 (Z(11)= 1.42, *p* = 0.382).

To further analyse these significant, attentionally-driven classification patterns, we performed pair-wise pattern classification to determine which attentional states generated different voxel-level responses. Significance was assessed using one-sample Wilcoxon signed-rank tests versus chance (50%), Benjamini-Hochberg corrected for the number of comparisons across ROIs.

In V1, we found above-chance classification of attention to orientation versus shape (Z(11)= 3.06, *p* = 0.028) and contrast versus shape (Z = 2.94, *p* = 0.028). Within both V3A/B and LO-1, classification accuracies were significantly above chance for orientation versus contrast (V3A/B: Z(11) = 2.63, *p* = 0.042, LO-1: Z(11) = 2.98, *p* = 0.028) and orientation versus shape conditions (V3A/B: Z(11) = 2.94, *p* = 0.028, LO-1: Z(11) = 2.83, *p* = 0.034). Within LO-1, the classification of contrast versus shape data was also significantly above chance (Z(11) = 2.67, *p* = 0.042). Within hV4, successful classification of orientation versus shape and contrast versus shape above change level was identified (Z(11) = 2.75, *p* = 0.037 and Z(11) = 2.99, *p* = 0.028 respectively). Within LO-2, classification of both orientation versus contrast and orientation versus shape above chance level was identified (Z(11) = 2.59, *p* = 00.043 and Z(11) = 2.93, *p* = 0.028 respectively) (see Figure.3C).

### 1.3.4 Multivariate Analysis: spatial pattern analysis

Previous reports have shown that some types of bottom-up stimuli generate patterns of retinotopically-biased responses within individual areas at relatively large spatial scales. For example, vertical and horizontal gratings drive voxels near the vertical and horizontal midline respectively (Tootell et al., 1998; Freeman et al., 2011). Other researchers however, suggest many early-visual voxels contain more complex tuning properties than predicted by coarse-scale biases such as radial bias or cardinal orientation selectivity, and these voxels with varying preferences are intermingled within V1 (e.g. Kay et al., 2008; Mannion et al., 2009; Kamitani and Sawahata, 2010; Pratte et al., 2016; Alink et al., 2017). We asked whether coarse-scale biases, or fine-grain patterns of voxel selectivity might be responsible for the top-down, attentionally-driven multivariate classification results that we found here. Subjects, might, for example, always attend preferentially to a particular part of visual space in order to solve different types of shape or orientation discrimination tasks.

To answer this question, we identified the voxels that were most informative for each type of classification decision and back-projected these into visual space. This allowed us to average spatial patterns of voxel preferences across observers. If all subjects used a common strategy (for example, attending to the vertical meridian) for a particular task, these averages would reveal a consistent non-zero response in this location. If, on the other hand, no changes in the large-scale pattern of responses was generated by attention, these maps would average to zero. Figure.4 shows mean values were computed across participants and thresholded (+/− 1.7 z-score, *p* < 0.05) to produce feature-specific attentional modulation maps. No significant patterns of attentional modulation are evident for any featural attentional focus across ROIs, and the 2° annulus of averaged attentional modulation revealed no clear peak of spatial attentional focus as a function of polar angle (see Figure 4).

**Figure.4:**
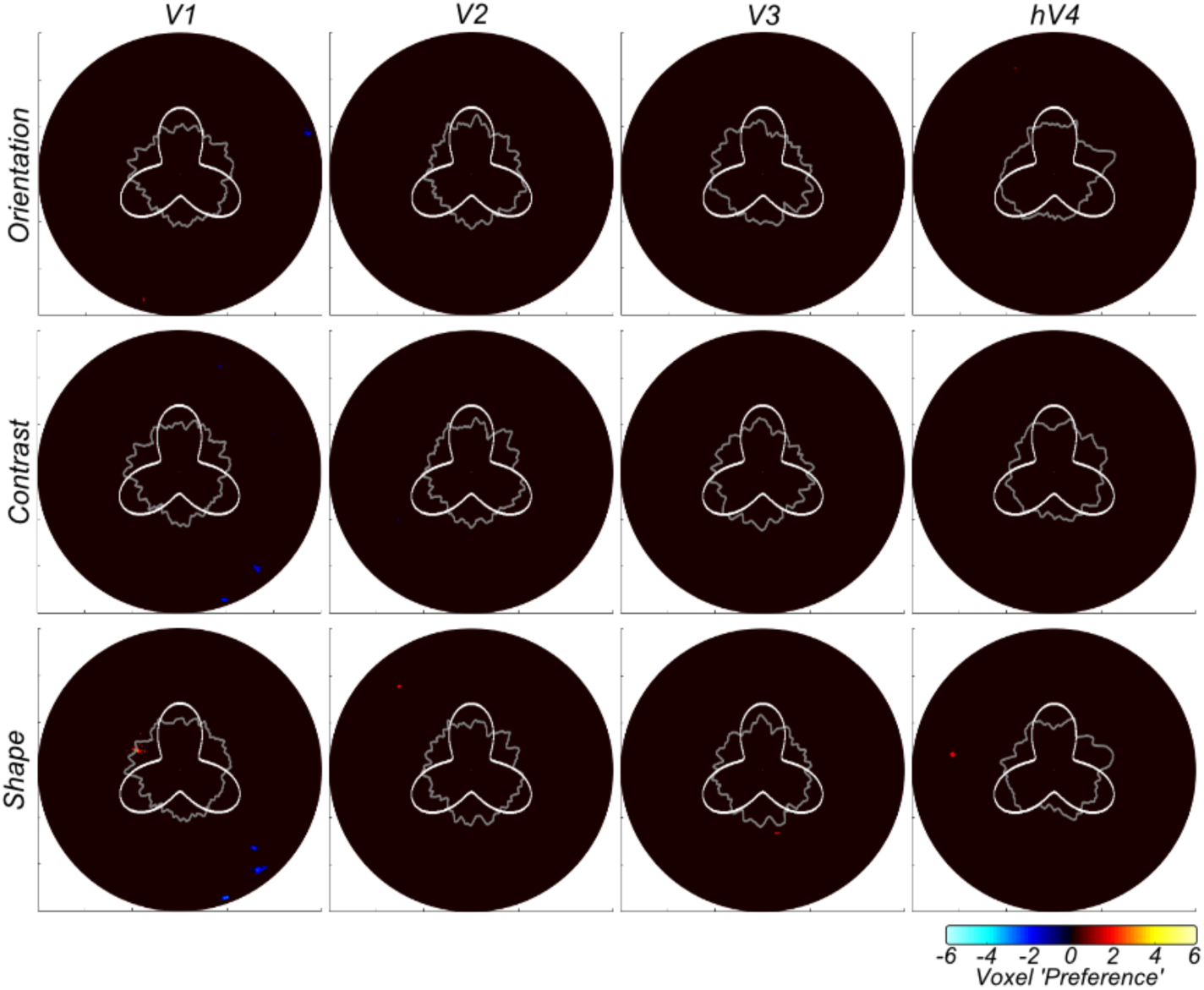
Group-averaged voxel feature-specific weights as a function of eccentricity (6°) and polar angle (360°) reveal no large-scale biases in voxel weights across location. The gray annulus reflects averaged voxel modulation at 1° intervals across 1.5-2.5° visual space. Deviations from circularity indicate positive (feature-specific) or negative (passive viewing) preferring clusters of voxels. A radial frequency pattern stimulus overlay is provided for reference. Activation is thresholded at +/− 1.7 z-score (p <0.05).

We conclude that while the stimulus modulations that we use to drive attentional tasks may be too subtle to drive different voxel-level BOLD responses, the different attentional states that subjects employ to detect these changes do select different neuronal populations within early visual areas. Our back-projection analysis indicates that these different populations are not consistent at a large-spatial scale across subjects. Our results are consistent with the hypothesis that subjects are selecting neurons from interdigitated populations that are optimal for particular tasks.

### 1.3.5 Timeseries Connectivity Analysis

Attention had relatively little effect on time-averaged univariate BOLD responses in individual visual areas. In our final analysis, we asked whether attention altered the way that individual ROIs communicate with each other. Specifically, asked whether functional connectivity (as measured by the similarity of timecourses in different ROIs) might be altered when subjects change their attentional state.

We first performed non-parametric Kendall’s Tau correlations between univariate (ROI-averaged) timecourses from each pairwise combination of visual ROIs (V1, V3A/B, hV4, LO-1, LO-2 and IPS0) for each featural attention condition. These inter-ROI correlation patterns represent a ‘fingerprint’ for each task (see Figure.5A). We asked if this overall fingerprint was altered by attentional task and then examined more detailed, pairwise combinations of attentional condition through a one-way repeated measure ANOVA with Bonferroni-corrected post-hoc tests, conducted on normalised and Fisher-z transformed coefficients.

**Figure.5:**
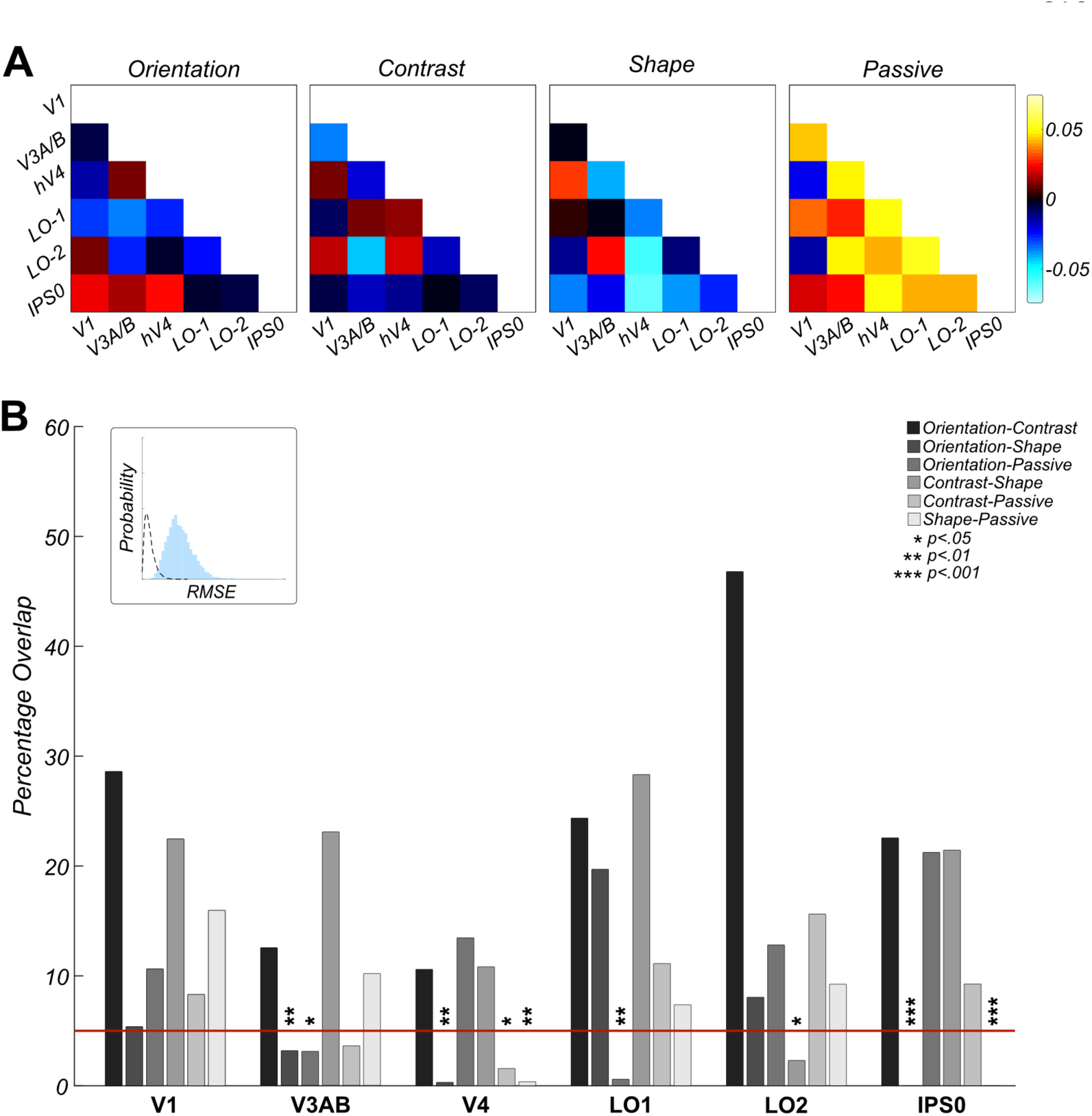
Greater overall connectivity during passive viewing compared to directed attention. A) Group feature-specific averaged attentional modulation connectivity values, indicating significantly greater mean connectivity during passive viewing than in the attentional tasks. B) Bootstrapped measures of distance between pairwise combinations of attentional focus conditions, for V1, V3A/B, hV4, LO-1, LO-2 and IPS0 ROIs. The figure demonstrates the percentage overlap between the distribution of RMSE scores across 10,000 iterations of randomly-selected samples of the observed data (blue) and scrambled correlation ‘noise’ data (dashed line) (see inset histogram for example distributions), for each ROI across multiple pairwise comparisons. A reference red line indicates 5% percentage overlap. Significant overlap (less than 5%) between pairwise combinations of condition are indicated with asterisks.

Most strikingly, our analysis revealed significantly greater positive correlation between ROIs during *passive* viewing than any attentional task condition (F(3,42) = 11.03, *p* <0.001, Bonferroni-corrected post-hoc tests, orientation versus passive; *p* = .003, contrast versus passive; *p* = 0.006, shape versus passive; *p* = 0.002). On average, there were no differences in the total level of connectivity between attentional conditions although, as shown below, individual pairs of areas do show significantly different levels of functional correlation in different attentional tasks.

To identify the specific ROI-ROI connectivity patterns driving these differences in correlation ‘fingerprint’ across attentional tasks, we calculated the Euclidian distance (RMSE) between pairwise comparisons of attentional task condition (averaged across a subsample of participants for each condition), for both observed and noise (scrambled attention-condition label) datasets, bootstrapped across 10,000 iterations.

Comparing patterns of connectivity associated with IPS0 (IPS0-V1, IPS0-V3A/B, IPS0-hV4, IPS0-LO-1, IPS0-LO-2) revealed a difference between orientation and shape attentional focus conditions significantly larger than expected by chance (*p* <.001). We also discovered significant differences in IPS0 connectivity between shape versus passive attentional focus conditions (*p* <0.001). hV4 connectivity across conditions revealed a similar pattern of results, with significant differences in connectivity between orientation and shape attention conditions (*p* = 0.003) and shape versus passive conditions (*p* = 0.006). There was an additional significant difference between activation patterns across hV4 connectivity between the contrast and passive conditions (*p* = 0.017). Analysis across LO-1-correlated ROIs revealed a significant difference in patterns of connectivity between orientation and passive conditions (*p* = 0.004) and we identified a significant difference between contrast and shape attentional focus conditions across LO-2-correlated ROIs (*p* = 0.022). However, connectivity to and from V1 (V1-V3A/B, V1-hV4, V1-LO-1, V1-LO-2, V1-IPS0), and V3A/B (V3A/B-V1, V3A/B -hV4, V3A/B -LO-1, V3A/B -LO-2, V3A/B - IPS0), did not appear to change as a function of task (*p* > 0.05) *(see Figure.5B)*.

## 1.4 Discussion

Here, we show that both *within*-area neuronal activity and *between-*area connectivity change as subjects switch between visual tasks. Perhaps surprisingly, at the univariate level we identify no differences in bottom-up BOLD signal modulation in response to differences in the visual features themselves: brief changes in orientation, contrast and shape all elicited essentially the same pattern of activation at the level of individual areas. Additionally, we identify very few areas that exhibit attentionally-driven changes in averaged BOLD signal, supporting the findings of previous attentional modulation literature (Sumner et al., 2008; Seymour et al., 2010; Song et al., 2011). This null finding is not due to neuronal response saturation: We see both robust bottom-up and top-down modulations in all visual regions in our data: areas are modulated by both subtle stimulus changes and differing attentional focus. However, these modulations are not dependent on the nature of the change. Reassuringly, we also identify no significant differences in univariate modulation between attention to orientation, contrast and shape in our control auditory cortex ROI (A1), suggesting any univariate differences we do identify are restricted to visually-responsive regions of cortex. Univariate changes in individual visual areas are therefore relatively uninformative about either bottom-up stimulus parameters or attentional state.

As an example, area LO-1 responds robustly to changes in all three stimulus parameters. This result is intriguing because visual areas are often characterised by their response to bottom-up changes in specific stimulus features and this area has been identified previously as having a causal role in orientation processing (Larsson and Heeger, 2006; Silson et al., 2013). Our data suggest that measuring univariate BOLD activity in visual cortex as a function of either stimulus type or task may not provide a complete picture of the computations that are being performed in each area. This finding is consistent with a model of visual cortex where areas typically contain multiple, overlapping feature maps in which activity depends both on the stimulus and, to an equal degree, on the attentional state. However, from univariate analysis alone, it is unclear whether there are simply no feature-specific attentional modulation effects, or whether by reducing ROI activity to a single number, potential fine-grain differences in feature-specific patterns of attentional modulation are lost.

To answer this question, we then analysed pattern changes at the voxel level. We showed that response *patterns* within individual visual areas are highly selective for the visual task and can, therefore, be used to decode attentional state. This supports the findings of previous decoding analyses (Kamitani and Tong, 2005, 2006; Sumner et al., 2008; Brouwer and Heeger, 2009; Freeman et al., 2011; Song et al., 2011).

Classification performance depends both on area and conditions: For example, in V1 successful classification was evident for all pairwise combinations of attentional modulation, supporting literature demonstrating the influence of attention at the earliest stage of the cortical hierarchy (Tootell et al., 1998; Serences and Boynton, 2007; Seymour et al., 2009, 2010; Lauritzen et al., 2010; Verghese et al., 2012). We believe these successful classifications reflect allocation of top-down attentional focus as oppose to bottom-up stimulus driven effects of selection history as suggested by Awh et al., (2012) and Theeuwes, (2013). Within our experiment, each attentional block was repeated the same amount of times, with the presentation order randomised across runs and participants. Additionally, changes in stimulus feature were randomised (within their 20% probability of change occurrence) hence stimulus information and task difficulty remained constant across blocks. Therefore, we believe our findings reflect voluntary allocation of top-down attentional focus, rather than any by-product of differential priming, arousal or selection history across conditions.

The more selective patterns of decoding performance within V3A/B and hV4 suggest that attentional exerts feature-specific effects along both the dorsal and ventral pathways of visual cortex. It clearly demonstrates that both streams possess neuronal populations with feature-specific preferences and that these neuronal populations are responsive to attentional demands (or else inherit attentionally-driven modulations from earlier in the visual pathway). Within LO-1, successful classification was evident across all pairwise combinations of visual task, but LO-2 exhibited successful decoding of orientation versus contrast and orientation versus shape only, partial support for the original conclusions of previous literature, with differing patterns of activation across orientation and shape attention (Larsson and Heeger, 2006; Silson et al., 2013). Within A1, we demonstrate no classification accuracies significantly greater than would be expected by chance, adding strength to our findings and demonstrating the specificity of our results within the visual cortex. Overall, these findings extend those from more traditional univariate analyses, and indicates the brain is able to up- and down-weight activity of specific neurons within particular visual regions in a task-dependent manner, in agreement with many previous reports (Treue and Martínez Trujillo, 1999; Martinez-Trujillo and Treue, 2004; Serences and Boynton, 2007; Serences et al., 2009; Verghese et al., 2012).

There is some debate in the literature about the origin of the information that drives fMRI multivariate analyses. Although the ability to decode at above chance rates from a visual area demonstrates that voxel activation patterns in that area depend on the experimental condition, the spatial scale of this change is important for interpreting the results. It has been shown that certain types of information are decoded from changes in voxel activity at a far coarser scale: In particular, researchers have demonstrated that grating orientation is encoded largely by retinotopically-driven, low spatial frequency patterns of response that switch from the vertical to the horizontal midline for vertical and horizontal gratings respectively (Freeman et al., 2011). Yet, if changes occur at the level of individual voxels, they may be driven by neuronal modulations at the level of columnar-scale tuning maps: for example, it has been hypothesised that the ability to decode local radial biases from primary visual cortex is driven by selective activation of orientation-selective neurons within the orientation pinwheels (Mannion et al., 2009). Additionally, recent work has provided evidence for the complexity of voxel tuning profiles within the early visual cortex and demonstrated that experimental task design can influence the conclusion that radial bias is the only source of orientation information within fMRI signals, for example (Pratte et al., 2016). Hence, global areal maps are unlikely to fully account for the ability to decode orientation signals within early visual cortex (e.g. Alink et al., 2017).

Although the stimuli used here varied only slightly in their physical characteristics, it is possible that MVPA performance was still driven by gross changes in attentional focus. Subjects might have adopted a spatially-driven strategy to solve different featural tasks – focusing on the very top of the radial frequency pattern, for example, to detect orientation changes. In Figure.4 we demonstrate that decoding does *not* depend solely on large-scale biases in sensitivity within visual areas. We find no evidence of a consistent spatial bias in the voxels used for different types of decoding. Although in principle, it is possible that such biases exist on a subject-by-subject (or even trial to trial) basis, our data are consistent with the hypothesis that attention selects sub-populations of relatively fine-scaled maps within individual visual areas. We suggest multivariate pattern classification shown here is derived from changes in activation at a relatively fine scale although we do not rule out coarser-scale topographically-determined responses that may vary across subjects ((Kamitani and Sawahata, 2010; Op de Beeck, 2010; Freeman et al., 2011; Pratte et al., 2016; Alink et al., 2017).

Our connectivity results show that visual processing is a dynamic, interactive process that is dependent on the task. While attentional effects have been noted in fMRI research since the late 1990s (Kastner et al., 1998; Tootell et al., 1998), investigating connectivity between regions of visual cortex as a function of task is a relatively novel approach.

Connectivity changes dramatically as a function of attentional state: Overall, we identify significantly greater average connectivity between ROIs during passive viewing than directed attentional focus. This enhanced overall connectivity during passive viewing is similar to the pattern of connectivity observed in the ‘default mode’ network (DMN) (e.g. Gusnard and Raichle, 2001; Raichle et al., 2001; Christoff et al., 2009), which is abolished by attentional task – and indeed the DMN includes parts of visual cortex. Once a visual task is provided, the brain switches to a more specific connectivity pattern and this pattern is task dependent. We identify different patterns of connectivity when subjects change attentional state for orientation, contrast and shape.

A previous fMRI experiment demonstrated that attention to a particular object category (faces or scenes) lead to strengthened coupling between category-selective visual areas (FFA and PPA respectively) and early visual cortex (e.g. Al-Aidroos et al., 2012). For example, they demonstrated attention to faces increased the proportion of intrinsic variance shared between regions of the occipital cortex and the FFA. In unpublished research, examining patterns of connectivity across the visual cortex during attention directed towards face stimuli, we also identify increased connectivity between visual areas in comparison with attention directed to low-level visual features. However, here, we identify *reduced* connectivity between visual areas during attention to low-level visual stimulus attributes in comparison to passive viewing. We believe this reflects the nature of our experimental design. In this experiment, participants directed attention towards highly controlled low-level stimulus attributes in a challenging task, in comparison to attention directed towards relatively higher level, complex features (faces and scenes) that are relatively independent of low-level cues. Here, we also have an explicit passive viewing condition for comparison. Therefore, it is likely we see differing effects of attention on functional connectivity within the visual cortex as a reflection on the type of stimulus attended and the task employed.

We examined these different connectivity patterns across ROIs as a function of task and identify different patterns of connectivity between certain ROIs for different visual tasks. Intriguingly, V1 is *not* one of those areas; correlations between V1 and other areas do not appear to change significantly depending on visual task (although they are reduced overall compared to the passive condition). However, we identify a difference in correlation between attentional tasks in IPS0, supporting a wealth of previous literature indicating the role of IPS in the modulation of top-down attention (e.g. Di Russo et al., 2003; Bressler et al., 2008; Lauritzen et al., 2009; Buffalo et al., 2010). Areas hV4, LO-1 and LO-2 have different patterns of connectivity across ROIs in directed attention. These findings indicate different fingerprints of attention across ROIs: not all ROIs and their connections are modulated the same way in all attentional tasks. Perhaps these regions selectively disengage or de-correlate with other networks during feature-specific directed attentional focus in order to process these features most effectively as previous cognitive flexibility research suggests (e.g. Spadone et al., 2015; Vatansever et al., 2016; Reineberg et al., 2018). Future research could seek to identify the link between specific patterns of attentional modulation across time within the visual cortex and indices of participant ability, through a network, as oppose to more traditional region-specific approach.

We believe our findings here are not a reflection of gross differences physiological between our four attentional conditions. For example, it is unlikely factors such as extraneous eye movements or changes in heart rate were responsible for differences we see in fine-grain voxel-level activation maps or patterns of functional connectivity. Our experiment recruited experienced observers, who are well-trained in maintaining a constant central fixation within 10-20 minutes of arc (Kowler, 1990). Additionally, our visual stimuli were present for 200ms, shorter than the time needed to make a visual saccade (Carpenter, 1988). Finally, if such gross-scale differences were apparent between attentional conditions, we would expect to see these differences evident at a univariate level. Instead, we identify relatively few significant univariate differences between our featural attention conditions and with passive viewing data, hence, our results likely reflect differences in attentional focus, rather than any consistent differences in gross-scale measures of arousal.

To conclude, we have used a relatively novel approach for investigating top-down attentional modulation signals in visual cortex. We show clear evidence of attentional modulation from the earliest stage of the visual cortical hierarchy and suggest that directed attention produces local voxel-level changes in activation as oppose to reflecting global topographical organisation of visual regions. Connectivity analyses demonstrate that attention causes a strong *decorrelation* of ROI responses relative to the passive state, which appears to be mediated by top-down signals processed within specific visual regions. This paradigm is a useful tool to probe the influence of a common confound within visual neuroscience, examining activation in response to shifting attentional focus rather than stimulus driven changes.

## Acknowledgements

We thank Theo Karapanagiotidis for his helpful contributions.

